# Shared Nearest Neighbor clustering in a Locality Sensitive Hashing framework

**DOI:** 10.1101/093898

**Authors:** Sawsan Kanj, Thomas Brüls, Stéphane Gazut

## Abstract

We present a new algorithm to cluster high dimensional sequence data, and its application to the field of metagenomics, which aims to reconstruct individual genomes from a mixture of genomes sampled from an environmental site, without any prior knowledge of reference data (genomes) or the shape of clusters. Such problems typically cannot be solved directly with classical approaches seeking to estimate the density of clusters, e.g., using the shared nearest neighbors rule, due to the prohibitive size of contemporary sequence datasets. We explore here a new method based on combining the shared nearest neighbor (SNN) rule with the concept of Locality Sensitive Hashing (LSH). The proposed method, called LSH-SNN, works by randomly splitting the input data into smaller-sized subsets (buckets) and, employing the shared nearest neighbor rule on each of these buckets. Links can be created among neighbors sharing a sufficient number of elements, hence allowing clusters to be grown from linked elements. LSH-SNN can scale up to larger datasets consisting of millions of sequences, while achieving high accuracy across a variety of sample sizes and complexities.

## 1 Introduction

Clustering is usually defined as the task of unsupervised learning, where the class labels of the data items are unknown [4, 21, 29, 35]. Clustering methods aim to create categories from the data in such a way that similar objects will be grouped together, while dissimilar objects will be separated into different groups, referred to as clusters. Important issues in clustering research focus on the effectiveness and scalability of the methods on data of varying complexities and arising from various domains [1, 39, 56].

Commonly used methods to cluster high dimensional data are presented in [37]. *K*-means is one of the most widely used clustering method due to its low algorithmic complexity. However, it has been shown in [53] that *K*-means tends to produce clusters of relatively uniform sizes and globular shapes, even if the data structure is endowed with varying cluster sizes or different shapes. This bias is known as the uniform effect of the *K*-means. Moreover, the number of clusters *K* has to be specified *a priori,* which is not trivial when no prior knowledge is available. To address these problems, methods based on estimating the density and/or the similarity among instances have been introduced [15, 30].

In [14], the authors presented an effective clustering method based on two key notions: the similarity between neighboring elements and the density around instances. This method, Shared Nearest Neighbors (SNN), is a density-based clustering method and incorporates a suitable similarity measure to cluster data. After finding the nearest neighbors of each element and computing the similarity between pairs of points, SNN identifies core points, eliminates noisy elements and builds clusters around the core elements. This method can yield better performance compared to other clustering approaches with data of varying densities, and it can automatically handle the number of output clusters. However, this method has complexity *O*(*n*^2^), where *n* is the number of instances in the dataset, arising from the computation of the similarity matrix, which can be prohibitive when dealing with high dimensional data.

One interesting concept to reduce the burden of computing the similarity matrix is Locality Sensitive Hashing (LSH). This concept was initially introduced to find approximate near neighbor information in high dimensional space [19, 51]. The key idea is to hash elements into different buckets; then for a query instance x, to use instances stored in buckets containing x as candidates for near neighbors. This approximation reduces the query time complexity to *O*(log *n*) instead of *O*(*n*) (*O*(*n*) is the complexity for searching nearest neighbors for one instance). Therefore, the similarity matrix computation time can be reduced to *O*(*n* log *n*).

We propose here to retain the basic principle of LSH by randomly splitting the dataset into a number of smaller-sized subsets, using a family of hashing functions, so that similar elements will be hashed together with high probability. We then look for nearest neighbors of each element in its bucket, and construct links among elements sharing a significant number of neighbors in order to output clusters. The proposed method, called LSH-SNN, has the advantage of reducing the complexity for computing the similarity matrix, while maintaining the same level of clustering accuracy.

In the present study, we have evaluated the performance of the LSH-SNN method on metagenomics datasets of various sizes and complexities. We also have compared the results with another density-based clustering method and the *K*-means method implemented in a popular sequence clustering software called MetaCluster [55]. Many computational tools have been proposed in the literature to analyze metagenomic sequences generated from micro-organism communities. These tools can be grouped into two main categories: (a) supervised and (b) unsupervised methods. Supervised methods, often relying on sequence similarities and alignments of DNA fragments to reference sequences of known taxonomic origins [59] include tools such as MEGAN [28] and CARMA [36]. Unsupervised methods group metagenomic fragments based on intrinsic features, such as the statistics of *l*-mer frequencies extracted from fragments. Unsupervised methods, such as MetaCluster [55], AbudanceBin [54] and TOSS [50], became attractive due to the lack of reference genomes for the bulk of micro organisms.

The rest of this paper is organized as follows: Section 2 surveys related work on clustering; Section 3 recalls some background on Local Sensitive Hashing and the Shared Nearest Neighbor methods; Section 4 introduces our method based on the combination of Local Sensitive Hashing and Shared Nearest Neighbors. Experimental results are illustrated in Section 5, while Section 6 concludes the paper.

## 2 Related work

Clustering methods look for similarities within a set of instances without any need for prior data labeling. Numerous methods have been proposed in literature to deal with clustering tasks. Existing algorithms can be grouped into five categories as proposed in [4] and [35]: partitioning methods, hierarchical methods, density-based methods, grid-based methods, and model-based methods. Hereafter, we will describe the main characteristics of these methods.

Partitioning methods construct *K* partitions of the data by grouping instances around the gravity center of each cluster. They can be divided into two main groups: the centroid methods such as *K*-means [20], and the medoids ones [33] such as the *K*-modes [25] and the *K*-prototypes algorithms [26]. Partitioning methods are simple to implement, however, the number of clusters *K* should be specified.

Hierarchical methods build a tree hierarchy, known as dendrogram, to form clusters in two different manners: agglomerative (bottom-up) and divisive (top-down). The former starts with singleton clusters and recursively merges them in a bottom-up strategy, while the latter breaks the dataset into smaller clusters in a top-down strategy. They use various local criteria to join or split clusters. Hierarchical methods have the advantage of handling any form of similarity without requiring the number of clusters to be known in advance. However, to construct a dendrogram, they suffer from their time and space complexities which are quadratic with respect to the number of clusters. Hierarchical clustering include methods such as: BIRCH [58], CURE [18] and CHAMELEON [31].

Density-based methods generate clusters based on the density of instances in a region. These methods are related to different concepts defining a point's nearest neighbors, such as density, connectivity, and boundary. Density-based methods are scalable and can find arbitrary shaped clusters; however, they output border instances, which may be unclustered and considered as outliers. Existing methods include DBSCAN [15], OPTICS [3], DENCLUE [23], Jarvis-Patrick [30], and SNN [13] algorithms. SNN will be described in further detail in Section 3.2.

Grid-based methods quantize the space into a finite number of cells that form a grid structure. Clustering is, then, performed on the grid cells, instead of the database itself [39]. The main advantage of these methods is their fast processing time; however, they output clusters with either vertical or horizontal boundaries. No diagonal boundary can be detected. This category includes STING [52], WaveCluster [48], and CLIQUE [2] algorithms, among others.

In model-based clustering, it is assumed that the data are generated from *K* probability distributions, and the goal is to find the distribution parameters [56]. Model-based methods are characterized by a small number of parameters; however, the computational burden can become significant if the number of distributions is large. Moreover, it is difficult to estimate the number of clusters. Many model-based clustering methods are described in the literature, such as Expectation-Maximization [41], SOM net [32] and AutoClass [10].

None of the above categories can directly cluster large amounts of instances of arbitrary shapes, and at the same time automatically detect the appropriate number of clusters. Shared nearest neighbors algorithm from the density-based clustering category can deal with local density variations and automatically find clusters of different shapes. However, adapting this technique with massive data requires extensive storage and time costs, especially for the step of computating the all-versus-all similarity matrix.

## 3 Background

In this section, we briefly review some background on local sensitive hashing (Subsection 3.1) and shared nearest neighbors algorithms (Subsection 3.2).

### 3.1 Local Sensitive Hashing

Local Sensitive Hashing (LSH) was first introduced in [19] as a classical geometric lemma on random projections, to quickly find similar items in large datasets. One or many families of hash functions map similar inputs to the same hash code. This hashing technique produces a splitting of the input space into many subspaces, called bins or buckets, with a high probability that instances originally close in their input space will be in the same bin or in adjacent bins within the LSH framework.

To alleviate the curse of dimensionality, each hash function projects the data to a lower-dimensional space (*h*: ℝ^*d*^ → ℤ). Different techniques have been presented in the literature to generate hash functions. These techniques can be categorized into two families: min-hash [7] and random projections. In document classification, min-hash is typically used when looking for textually similar documents by processing items and generating integers from strings of characters [38]. Random projections, on the other hand, are obtained via simple probability distributions like p-stable distribution [12], and sign-random-projection [9].

Dealing with large datasets, LSH is usually used with the nearest neighbors techniques [8] or for clustering data [6, 7]. To perform k-nearest neighbors, buckets, and sometimes their adjacent buckets [40], containing the query element are checked and all the existing instances are ranked according to their distances to the query element. To cluster high-dimensional data [22, 34], similar elements contained in the same bucket can be joined to output clusters in a hierarchical way [45].

### 3.2 Shared Nearest Neighbor

SNN (Shared Nearest Neighbor) is a density based clustering approach for finding groups of documents with a strong, coherent topic or theme [4], [14], [17],[42], [43], [49]. SNN handles clusters of widely differing sizes, densities, shapes, and having large amounts of noise and outliers. To exploit space density of the data, SNN uses the concept of similarity based on the shared nearest neighbor approach. The similarity matrix is sparsifyed by keeping only the k-most nearest neighbors (*knn*). The shared nearest neighbor graph is then constructed by creating links between pairs of instances having each other in their respective *knn* lists. The weight of the link can be calculated either as the number of shared neighbors between two *knn* lists or using the ordering of these shared neighbors.

The algorithm determines the type of each instance (core, border or noisy) by calculating its connectivity; i.e., the number of links coming out of this instance, which will be compared to *noisy* and *topic* thresholds. Noisy instances are discarded and will never be used in the clustering process. Core instances form final clusters with their connected elements. This algorithm is configured by means of four parameters, namely: the number of nearest neighbors, noted as *knn* hereafter, the *topic* threshold, and two other thresholds to add elements to clusters. Depending on the user-defined parameters, many of the border instances remain unlabeled because they are not connected to core elements.

## 4 Our proposal: the LSH-SNN algorithm

In this section, we describe the key idea of our algorithm, called LSH-SNN, and described in Section 4.1. We then describe how to tune the different parameters in Section 4.2.

### 4.1 Method description

Shared nearest neighbor (SNN) is a relatively effective unsupervised method to automatically find clusters of different shapes and densities. However, it is challenged by scalability issues arising on large datasets. For *n* data items and *knn* nearest neighbors, the computational complexity of SNN is *O*(*n*^2^), whereas its space complexity is *O*(*knn* ⁎ *n*). For large number of instances *n*, SNN can suffer from important scalability problems.

Our goal is to adapt a suitable framework for clustering large number of instances using SNN principles. It is motivated by large metagenomic datasets incurring high computational costs. To help reduce these costs, we consider the rationale of local sensitive hashing as a framework for the development of the SNN method. In this framework, the *n* data instances are randomly partitioned into a number of smaller-sized subsets called buckets, and for each fragment, we locate its approximate nearest neighbors inside its bucket. This approach, called LSH-SNN, has an advantage over SNN since it restricts the calculation of distances for each single fragment inside its bucket, whereas SNN needs to calculate *n* distance measures for each fragment before selecting the list of the nearest neighbors.

LSH-SNN begins with the extraction of features from the sequence fragments by computing the frequencies of all possible *l*-mers (substrings of length *l*) in each of them (Section 4.1.1). Nearest neighbors of all sequences are then computed by applying the LSH technique, which involves splitting sequences into different buckets in such a way that similar sequences end up in the same bucket with higher probability (Section 4.1.2). Elements stored in buckets containing a given sequence x are retrieved and ranked according to their distances to x in order to compute its nearest neighbors list. Shared neighbors are linked according to the SNN rule, and connected sequences form output clusters (Section 4.1.3). Finally, in the case of unclustered fragments, a last step is performed in order to assign them to the cluster most similar in terms of l-mer distribution (Section 4.1.4).

#### 4.1.1 *l*-mer frequency calculation

Sequence similarities are typically identified by comparing occurence patterns of relatively short DNA substrings of length *l* between the sequences [50, 55]. Two broad scenarii can be used to assess *l*-mer-based similarities: *abundance-based methods* make use of relatively large *l* values (*l* ≥ 20) in order to ensure the uniqueness of most *l*-mers [50], while *composition-based methods* rely on smaller *l* values. Since DNA is a combination of four different types of nucleotides (A,T,G,C), there are at most 4^*l*^ *l*-mer combinations forming the feature vector. The frequency of each *l*-mer combination is normalized by dividing the number of occurrences by the fragment length.

Because of nucleotide base complementarities, the size of the feature vector can be reduced by half, i.e., for a DNA sequence x of length *s*, the feature vector is given by:

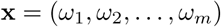

where 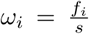, *f_i_* is the frequency occurrence of a given *l*-mer combination and *m* represents the size of the vector (or number of descriptive attributes), *m* = 4^*l*^ /2 if *l* is odd, and (4^*l*^ +4^*l*/2^)/2 if even.

#### 4.1.2 LSH

For convenience, we briefly recall some notations. Let 𝕏 be the collection of *n* sequences of *m*-dimensional features, and let x ∊ 𝕏 denote an input sequence. Let *k* be the number of projections. For each *i* ∈ [*k*], *h_i_*(x) is given by: *h_i_*(x) = sign(x.*v_i_*), where *v_i_* is vector whose components are randomly generated from a Gaussian distribution, for example 𝓝(0,1). This scalar projection gives one hash value for x. The hash code for x is then obtained by a concatenation of the *k* hash values, *g*(*x*) = (*h*_1_(x), *h*_2_(x),…, *h_k_*(x)). LSH prepares *r* copies of *g*(.) to improve the hashing discriminative power [11, 51] (to avoid confusion, *k* (in lower case) is the number of sampled bits, while *K* (in upper case) is the number of output clusters).

The feature vectors are first normalized with zero mean and unit variance. Each input sequence x is then indexed by a hash code *g*(x) = (*h*_1_(x), *h*_2_(x),…, *h_k_*(x)) defining its bucket identity, and the hash code of all sequences in 𝕏 constitutes a hash table. This projection produces a new *k*-dimensional space (*k* << *m*). Since the number of elements per bucket is typically much smaller than *n*, we need to ensure that similar sequences share the same bucket with higher probability while minimizing random effects. To achieve this, *r* hash functions *g*_1_,*g*_2_,…,*g_r_* are sampled independently, each generating a distinct hash table. For each sequence x, we then identify *g*_1_(x), *g*_2_(x),…, *g_r_* (x) indexing the *r* buckets where x mapped in each projection.

Note that any hash function may be applied in this step [44] when distances are measured as angles between point pairs. In this work, we demonstrate results based on the random hash functions generated from a Gaussian distribution.

The projection of the data items into different buckets can be summarized as follows:

##### Algorithm 1 Computation of *hash function*

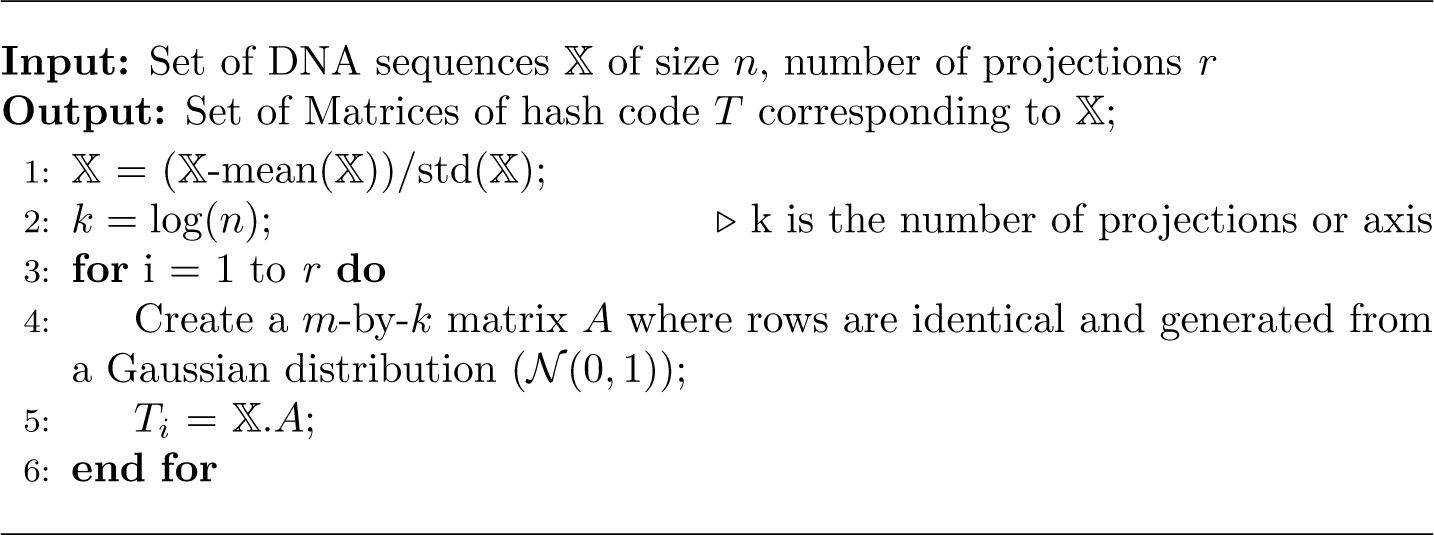

For hashing the dataset, the time complexity is *O*(*m* × *k* × *r*) per sequence, since each sequence of dimension *m* will be processed by *k* hash functions repeated *r* times. Therefore for *n* sequences the time complexity of LSH is *O*(*n* × *m* × *k* × *r*). For fixed *l*-mer (and hence of *m*) and *r* values, LSH has a complexity of *O*( *n* × log(*n*)).

#### 4.1.3 SNN

Once the space has been partitioned *k* × *r* times, a simple *k*-nearest neighbor classifier may be considered to find the nearest neighbors of a sequence x inside its bucket (i.e., having the same hash code) for all partitions. The union of *r* subsets of nearest neighbors for a given sequence is treated as its neighborhood list. Since the nearest neighbor lists are generated from sparsely populated buckets, the computational cost and runtime are improved.

Once the sets of nearest neighbors have been defined, the SNN algorithm follows two steps: computing link strengths and sequence labeling. A link is created between two sequences x_1_ and x_2_ if they have each other in their respective neighborhood lists, and it can be scored according to the sum of positions of shared instances between these two lists, namely:

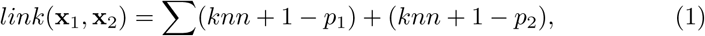

where *p*_1_ and *p_2_* are the positions of a shared neighbor in the lists of x_1_ and x_2_. The *knn* lists are then transformed into a graph where sequences (nodes) are connected via link strengths. For each sequence x in the graph, the sum of its total links *(conn_x_)* is computed in order to enable the selection of a subset of representative sequences according to a connectivity-based criterion (*conn > topic threshold*).

The algorithm 2 inset summarizes the SNN method. To check the nearest neighbors of a sequence x, *n*′ distances need to be evaluated, where *n*′ is the number of elements sharing the same hash code as x. Since we have *n* sequences, the time complexity for computing the nearest neighbor elements is *O*(*n* × *n*′ × *m* + *C*(*knn*)), where *C*(*knn*) is a relatively small factor enabling the selection of *knn* near neighbors for each sequence [30]. To estimate the link between two sequences having each other in their respective *knn* list, two columns of size *knn* are selected and evaluated. The cost of this process amounts to *O*(*n* × *knn* × *knn*). Therefore, the total complexity of the algorithm becomes *O*(*n* × *m* × *k* × *r* + *n* × *m* × *n*′ + *n* × *knn* × *knn*).

This algorithm is able to handle clusters of different densities. However, it can leave a large number of non-noisy sequences unclustered. To alleviate this problem, we define a new step to relabel unclustered sequences.

#### 4.1.4 Relabeling

A relabeling step was thus developed to reduce the number of unclustered sequences. It identifies a subset of frequencies characteristic of each cluster and contributing most to the classifier's accuracy, and discards other less relevant features. Each unclustered sequence is then added to the cluster most closely related with respect to the subset of frequencies.

Relabeling proceeds by computing the mean of each cluster and dividing it by the mean of the other clusters. Discriminant *l*-mer frequencies; i.e., those which most differentiate this cluster from others, are selected. For a given sequence, we compute its distance to the mean of each cluster by using the subset of discriminant frequencies and assign it to the nearest cluster. If two clusters are almost equally close to a given sequence, we keep the latter unlabeled in order to avoid increasing the number of misclassified instances.

The implementation of this part of the algorithm is presented as pseudo code in the algorithm 3 inset.

##### Algorithm 2 Computation of *SNN*

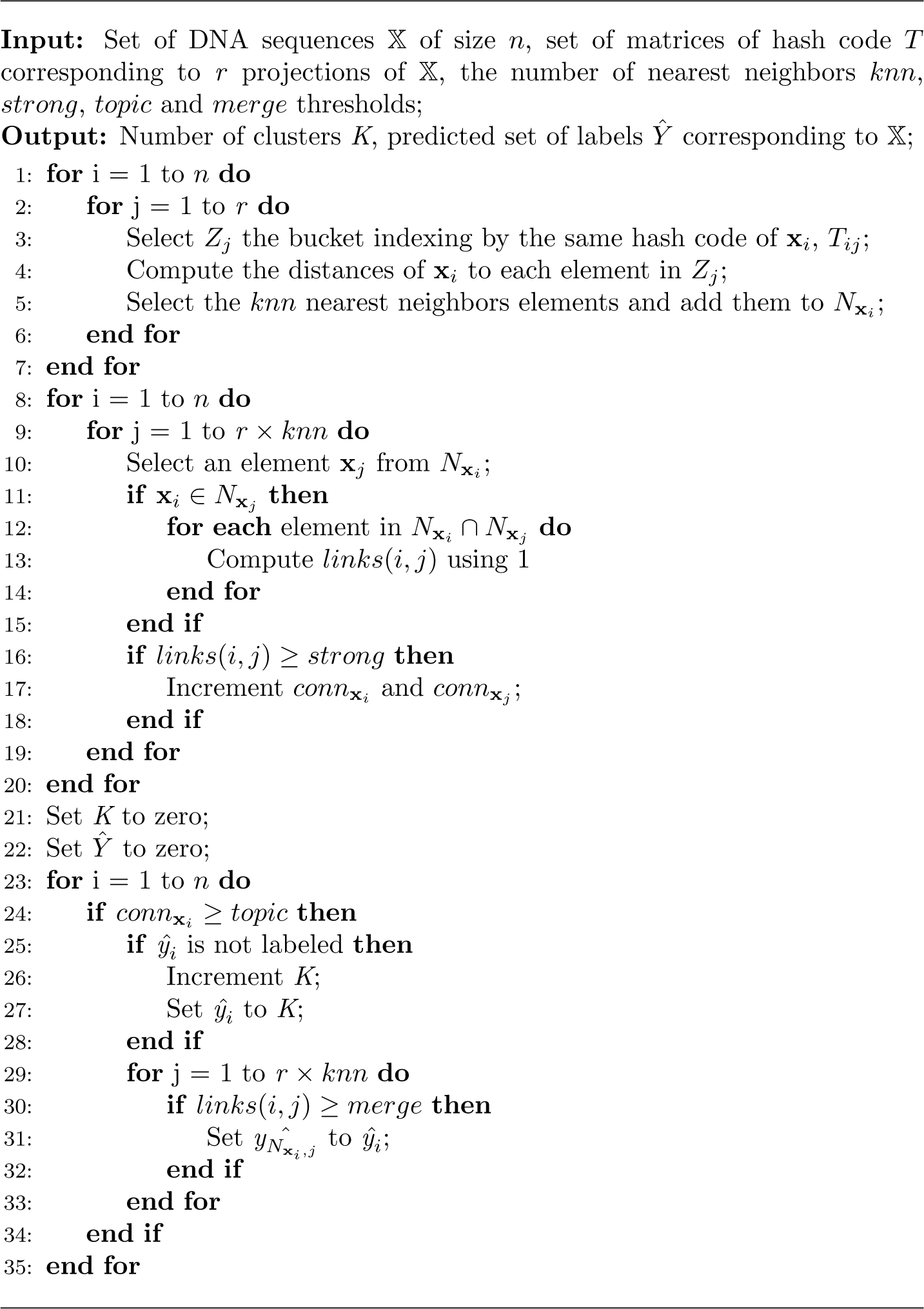

##### Algorithm 3 Relabeling of unclustered sequences

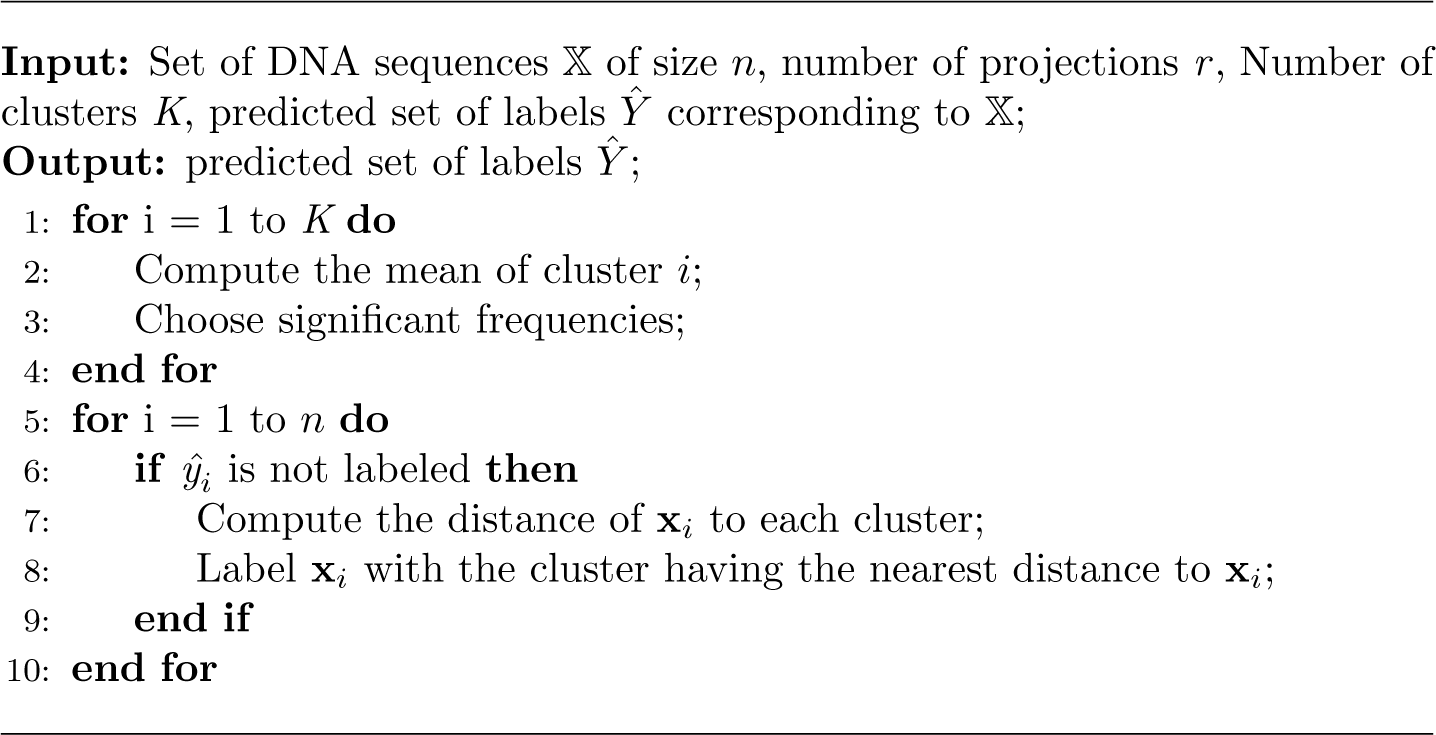

Relabeling requires *n* × *K* operations, hence a complexity of *O*(*n* × *K*). The overall complexity of the algorithm depends on the complexity of the hashing functions and the SNN classifier used. Since *l*-mer and *r* have fixed values for all experiments, the total complexity is *O*(*n* × log(*n*)+ *n* × *n*′ + *n* × log(*n*′) × log(*n*′)).

### 4.2 Parameters in LSH-SNN

This section discusses the configuration of the LSH and SNN parameters, which impact the method's performance both in terms of runtime and clustering quality. Parameters were determined by grid search and focused on optimizing the V-measure (see Section 5.3).

LSH has two parameters, *k* and *r*, to be tuned: (a) The number of sampled bits *k* determines the number of instances inside the buckets, which on average is expected to be equal to 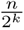. The total number of buckets is limited to max(*n*, 2^*k*^). If *k* takes a small value with respect to the number of sequences, then we would end up with a large number of sequences per bucket and the time-consumption of the SNN part will be very high. On the other hand, if *k* = *n*, we would get on average one sequence per bucket and there will be no knn lists to be constructed. In the present study, we set *k* to log(*n*). (b) The number of projections *r* is the second parameter. For *r* = 1, two close elements could end up in distinct buckets because of the random nature of the hashing. By increasing the number of projections, we increase the probability that these two elements are mapped to the same bucket in at least one projection. The number of projections *r* should thus be increased to stabilize the results. On the other hand, for large values of *r*, distant elements may be mapped to the same bucket, provoking the *r* × *knn* lists to grow and ultimately leading to the same drawbacks as the initial SNN method. In the present study, *L* was empirically set between 300 and 1200, depending on the size of the dataset.

Regarding SNN, four parameters influence the outcomes: *knn, topic, merge* and *strong* thresholds. (a) The size of the near neighbor list *knn* depends on the number of elements per bucket. To construct *knn* lists containing a sufficiently large number of closely related sequences consistent with the shared neighbors criterion, we need to choose a large number of nearest neighbors. However, increasing *knn* may add more distant sequences to the same cluster, thus increasing the computational cost. Decreasing *knn,* on the other hand will result in many smaller-sized clusters. A simple and pragmatic approach is to set 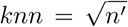 or *knn* = log(*n*′), where 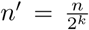. In our experiments, we fixed *knn* to log(*n*′). (b) The *topic* threshold determines the proportion of most highly connected links (i.e., having highest *connectivity)* to be selected as representatives. This threshold ranged from 0.04 to 0.06 in our experiments. (c) The *merge* threshold represents the percentage of links to be used in the cluster merging process, and was fixed to 0.02 in our experiments. (d) The *strong* threshold is used to reduce the number of unlabeled sequences (singleton clusters) and to choose representative elements. In our experiments, we set this parameter to 0.1.

## 5 5 Experiments

In this section, we detail the experimental setup. We first describe the datasets (Section 5.1) and provide a short reminder about the methods we compare our algorithm with (Section 5.2). We then detail the metrics used to evaluate the performance of our method (Section 5.3), and finally present and discuss the results (Section 5.4).

### 5.1 Datasets

We have used synthetic datasets of increasing sizes and complexities composed of 600 base-pair length reads (DNA fragments). The mean coverage of the datasets was fixed to 1X and 10X for two distinct series, which means that, on average, a given position in the genome is covered by 1 or 10 different reads respectively. The number of reads derived from each species is equal to 5 000 for 1X datasets and to 50 000 reads for 10X datasets [16].

To evaluate the performance of the various clustering modules on the benchmark datasets, we compared class memberships of elements (reads) in each dataset to the memberships induced by the clustering. Class membership of elements is trivial to define for datasets used in the composition-based clustering experiments, simply consisting in the genomes the read were sampled from, and the cardinality of the class set matching the samples richness. For all the synthetic datasets, the read generation process was performed using the mason software [24] with default error model parameters for Illumina reads (mason can insert position specific sequence modifications according to empirically calibrated and sequencing platform dependent error models).

### 5.2 Benchmark methods

The proposed method is compared with MetaCluster [55], a popular compositional binning software based on the K-means algorithm with the Spearman footrule distance, which operates on relative rankings of the *l*-mer frequencies [55].

The main advantage of this approach is its simplicity, which underlies its ability to handle datasets featuring a relatively large number of species. However, its behavior is sensitive to the random choice of initial cluster centers, and it may fail to output clusters when data are of non-globular shapes. Moreover, the number of clusters should be specified by users, which is not trivial when no prior knowledge is available [46].

The complexity of the *K*-means method is *O*(*n* × *m* × *K* × *Times*), where *n* is the number of instances, *m* is the dimension of data, *K* is the number of clusters and *Times* is the number of iterations for convergence. As *l*-mer is fixed for all experiments, the overall complexity of MetaCluster becomes *O*(*n* × *K* × *Times*).

We also compare our algorithm to the Jarvis-Patrick method (JP) [30], combined with the LSH indexing in a way similar to the LSH-SNN coupling previously described. JP also relies on the near neighbors similarity concept, but simply merges elements in the same cluster if they have a sufficient number of shared neighbors, i.e., the number of shared elements between two neighbors is greater than a user-predefined threshold *kt,* which we fixed to (*r* ⁎ *kpp*)^2^/2. The complexity of LSH-JP is *O*(*n* × *k* × *r* + *n* × *n*′ + *n* × *kpp* × *kpp*).

### 5.3 Performance evaluation

To evaluate the performance of our method, we considered the following metrics described in the literature: Homogeneity, Completeness, V-measure, F-measure and the Adjusted Rand Index.

*Homogeneity* evaluates the class distribution within each cluster. It is high when each cluster contains only elements of a single class. *Completeness,* on the other hand, examines the distribution of cluster assignments within each class. It is high when elements of a single class are assigned to a single cluster. Let *C* be the number of species in a dataset of *n* sequences and *K* be the number of output clusters. These two measures are given by [47]:

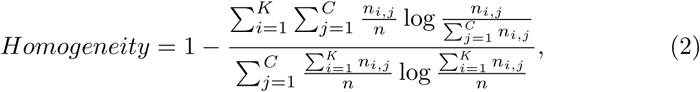

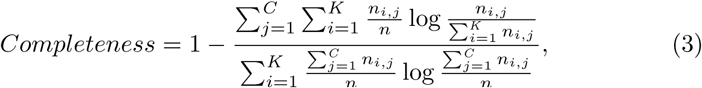

The V-measure is defined as the harmonic mean of *homogeneity* and *completeness*. It is calculated as:

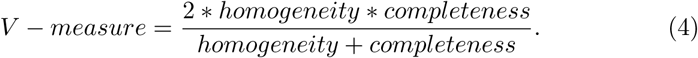

The clustering accuracy, *F-measure*, is defined as:

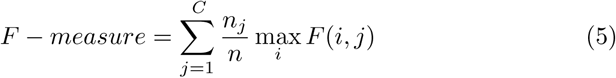

and,

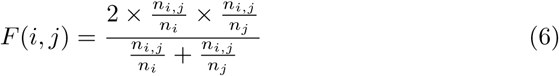

where *n_j_* is the cardinality of cluster *C_j_*.

*The Adjusted Rand Index* [27], [57] computes a similarity measure between the computed and the ideal clusterings as:

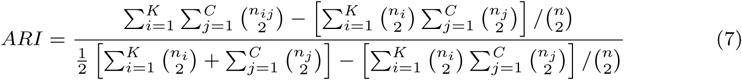

These measures take values between 0 and 1; higher values correspond to better clustering accuracy.

### 5.4 Discussion

After tuning the parameters described in Section 4.2, we evaluated the performance of LSH-SNN as well as two other clustering algorithms, LSH-JP and K-means (MetaCluster) on the different datasets (note that MetaCluster requires the number of component genomes to be given as input for each dataset). Clustering accuracy was quantified using different measures, shown in Table 1 which also displays the rank of each method and highlights (in bold letters) the best value for each evaluation criterion. The percentage of unclustered sequences (singleton elements) varied between 20% and 30% for LSH-SNN, 60% and 90% for LSH-JP, and between 2% and 4% for MetaCluster. These figures underly the lower Adjusted Rand Index and F-measure values achieved by LSH-JP. On the other hand, the relabeling step in the LSH-SNN algorithm specifically aims at reducing the number of unclustered sequences.

**Table 1:** Performance on synthetic datasets.

| Datasets | Metrics | LSH-SNN | LSH-JP | MetaCluster |
| --- | --- | --- | --- | --- |
| MC5-600-1X | Homogeneity | 0.502(2) | **0.614**(1) | 0.302(3) |
|  | Completeness | 0.504(2) | 0.499(3) | **0.618**(1) |
|  | V-measure | 0.503(2) | **0.551**(1) | 0.406(3) |
|  | F-measure | **0.642**(1) | 0.469(3) | 0.574(2) |
|  | Adjusted Rand Index | **0.645**(1) | 0.296(3) | 0.405(2) |
| MC10-600-1X | Homogeneity | **0.512**(1) | 0.499(2) | 0.415(3) |
|  | Completeness | **0.721**(1) | 0.704(2) | 0.632(3) |
|  | V-measure | **0.598**(1) | 0.584(2) | 0.501(3) |
|  | F-measure | **0.561**(1) | 0.461(3) | 0.522(2) |
|  | Adjusted Rand Index | **0.504**(1) | 0.312(3) | 0.417(2) |
| MC25-600-1X | Homogeneity | **0.516**(1) | 0.335(3) | 0.443(2) |
|  | Completeness | 0.548(3) | **0.629**(1) | 0.614(2) |
|  | V-measure | **0.531**(1) | 0.437(3) | 0.515(2) |
|  | F-measure | 0.320(2) | 0.304(3) | **0.476**(1) |
|  | Adjusted Rand Index | **0.349**(1) | 0.178(3) | 0.244(2) |
| MC50-600-1X | Homogeneity | **0.492**(1) | 0.353(3) | 0.389(2) |
|  | Completeness | 0.674(2) | **0.713**(1) | 0.594(3) |
|  | V-measure | **0.569**(1) | 0.471(2) | 0.471(2) |
|  | F-measure | 0.309(2) | 0.253(3) | **0.372**(1) |
|  | Adjusted Rand Index | **0.332**(1) | 0.141(3) | 0.167(2) |
| MC100-600-1X | Homogeneity | **0.492**(1) | 0.176(3) | 0.249(2) |
|  | Completeness | 0.674(2) | **0.704**(1) | 0.567(3) |
|  | V-measure | **0.569**(1) | 0.282(2) | 0.346(2) |
|  | F-measure | 0.309(2) | 0.141(3) | **0.227**(1) |
|  | Adjusted Rand Index | **0.332**(1) | 0.085(2) | 0.018(3) |
| MC5-600-10X | Homogeneity | **0.437**(1) | 0.269(3) | 0.297(2) |
|  | Completeness | 0.609(2) | 0.136(3) | **0.617**(1) |
|  | V-measure | **0.508**(1) | 0.181(3) | 0.402(2) |
|  | F-measure | **0.656**(1) | 0.032(3) | 0.573(2) |
|  | Adjusted Rand Index | **0.393**(1) | 0.003(3) | 0.144(2) |
| MC10-600-10X | Homogeneity | 0.421(2) | **0.779**(1) | 0.405(3) |
|  | Completeness | **0.666**(1) | 0.586(2) | 0.422(3) |
|  | V-measure | 0.516(2) | **0.669**(1) | 0.413(3) |
|  | F-measure | **0.457**(1) | 0.196(3) | 0.379(2) |
|  | Adjusted Rand Index | **0.364**(1) | 0.003(3) | 0.315(2) |

The name of the datasets are MCx-y-zX where x corresponds to the number of species, y is the read length and z indicates the mean coverage.

It can be seen that the behavior of the three algorithms remains almost the same on the MC5-600-1X and MC5-600-10X datasets, which is to be expected as these were generated from the same species with just ten times redundancy for the second dataset.

Table 1 illustrates that LSH-SNN outperforms LSH-JP and MetaCluster in terms of the *homogeneity* and *V-measure* metrics, the latter being the harmonic mean between *homogeneity* and *completeness*. LSH-SNN also slightly outperforms the other methods on the *F-measure* in four out of seven datasets, and consistently yields the best performance on all the datasets in terms of the *Adjusted Rand Index* metric. The latter result suggests that LSH-SNN has improved clustering accuracy on the datasets analyzed.

The LSH-SNN algorithm is implemented in the C++ programming language, and uses the OpenMP application programming interface to support multiprocessing. Experiments were conducted on a Linux x86_64 server endowed with multi-core CPUs and 2 TB of RAM. The LSH-SNN and LSH-JP computations were parallelized on 48 cores, while MetaCluster execution (which requires the number of clusters to be specified as an input parameter) distributed the computation across different threads according to the number of target clusters.

Overall, these results demonstrate that LSH-SNN achieves accurate binning for DNA sequences as short as 600 bp, as compared to LSH-JP and MetaCluster and despite the latter using the correct number of clusters (genomes) as an input parameter.

## 6 Conclusion

We have proposed an unsupervised composition-based method for binning large volumes of sequences, without any prior knowledge of their reference genomes or the number of distinct genotypes present in the analyzed sample. LSH-SNN is based on two essential steps: the hashing/indexing of the data space and the creation of links between sequences in order to output clusters. After computing the l-mer distribution of each sequence, LSH partitions the input space into buckets containing smaller subsets of sequences whose connectivity is evaluated based on the SNN rule. A third step was added to reduce the number of singletons or unclustered sequences.

The LSH-SNN algorithm can scale to datasets containing millions of sequences and does not require the number of output clusters to be predetermined. While the presented algorithm makes use of the SNN rule, we envision that the LSH concept could be combined with other clustering methods facing large data volumes, or used on its own as exemplified in [5], where a MinHash LSH scheme was used to compute similarities between long noisy reads generated with a new single-molecule real-time (SMRT) sequencing technology.

The LSH-SNN algorithm was evaluated on seven synthetic metagenomic datasets of different sizes and complexities (i.e., harbouring different numbers of organisms / genotypes). We observed that LSH-SNN performs comparably or better on these datasets than the two other clustering algorithms tested (LSH-JP and MetaCluster). We should note however that, even though LSH-SNN significantly increases the size of the datasets that can be handled as compared to what can be achieved with the SNN method alone, its complexity does not compare favourably with Lloyd's heuristic underlying most K-means clustering engines. Therefore, the latter is probably more suited to the analysis of larger datasets containing billions of sequences, which are already generated nowadays from complex metagenomics samples (e.g., from soil). Alternatively, the LSH-SNN approach could be applied to cluster the contigs (sets of overlapping sequences) resulting from a preliminary (meta)genome assembly step, instead of being applied to raw (unassembled) reads.

## Acknowledgments

This work was supported by the French Alternative Energies and Atomic Energy Commission (Commissariat l’Energie Atomique et aux Energies Alternatives), through its transversal programme Technologies pour la Santé (MetaTarget project). The authors would like to thank Anestis Gkanogiannis for generating and sharing the datasets, and Marcel Salanoubat for valuable discussions and overall support.

